# Annotating CryoET Volumes: A Machine Learning Challenge

**DOI:** 10.1101/2024.11.04.621686

**Authors:** Ariana Peck, Yue Yu, Jonathan Schwartz, Anchi Cheng, Utz Heinrich Ermel, Saugat Kandel, Dari Kimanius, Elizabeth Montabana, Daniel Serwas, Hannah Siems, Feng Wang, Zhuowen Zhao, Shawn Zheng, Matthias Haury, David Agard, Clinton Potter, Bridget Carragher, Kyle Harrington, Mohammadreza Paraan

**Affiliations:** Chan Zuckerberg Institute for Advanced Biological Imaging (CZ Imaging Institute), 3400 Bridge Parkway, Redwood City CA 94065, USA; Department of Biochemistry & Biophysics and the Howard Hughes Medical Institute, University of California, San Francisco, San Francisco, CA 94143, United States

## Abstract

Cryo-electron tomography (cryoET) has emerged as a powerful structural biology tool for understanding protein complexes in their native cellular environments. Presently, 3D volumes of cellular environments can be acquired in the thousands in a few days where each volume provides a rich and complex cellular landscape. Despite numerous innovations, localizing and identifying the vast majority of protein species in these volumes remains prohibitively difficult. Machine learning based methods provide an opportunity to automate the process of labeling and annotating cryoET volumes. Due to current bottlenecks in the annotation process, and a lack of large standardized datasets, training datasets for machine learning algorithms have been scarce. Here, we present a defined “phantom” sample, along with “ground truth” annotations, that will be the basis of a machine learning challenge to bring cryoET and ML experts together and spur creativity to address this annotation problem. We have also set up a cryoET data portal that provides additional diverse sets of annotated 3D volumes from cryoET experts across the world for the machine learning challenge.

## Introduction

Much of cell biology is still uncharted territory. The structural biology method of cryo electron tomography (cryoET) is uniquely poised to expand our understanding of cellular function in health and disease^1^. Samples are cryopreserved rather than chemically fixed for cryoET, which maintains structural integrity of cellular components, enabling us to peer inside cells and visualize molecular interactions with up to subnanometer resolution. When this technique is applied to lamellae^2,3^–frozen slices of cells–it provides a detailed 3D view (https://tinyurl.com/yx2t9wb9) of the cellular state at the moment of freezing^4^. In eukaryotic cells, this view includes many thousands of different protein complexes (tens of nanometers in size) and organelles (hundreds of nanometers in size)^5–7^.

Like tomography methods used in medical imaging such as Computed Tomography (CT) and Magnetic Resonance Imaging (MRI), cryoET also illuminates a region of interest (ROI) from different orientations and collects corresponding 2D projection images (Fig. 1a). These 2D images are then aligned and reconstructed to compute a 3D volume of the ROI (Fig. 1b), the result of which is called a tomogram. This type of data provides both the fine sampling required to obtain near-atomic structures of protein complexes (3-4 Ångstroms)^8,9^, and also the field of view (600-1000 nm) to visualize each protein complex in its biological context.

**Figure 1.**
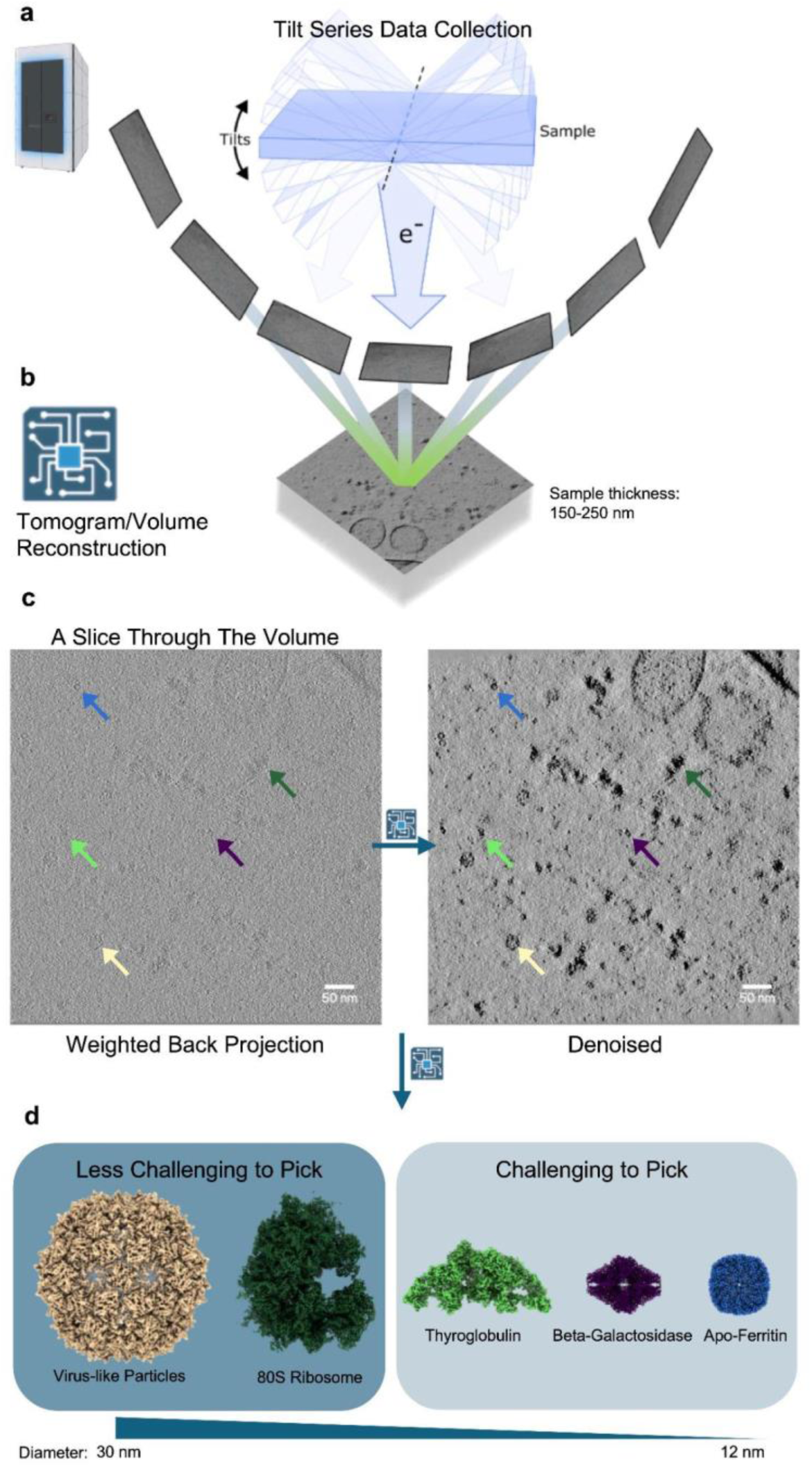
Cryo-electron tomography and its challenges. **a**. Tilt series data collection inside a Krios G4 transmission electron microscope. The frozen sample (blue slab) is rotated to generate 2D projection images on a fixed direct electron detector. The 2D images have an SNR of ∼0.1. The range of the tilted images is only representative, typically there are 31 to 61 tilted images. **b**. A 3D volume is computed using real space back-projection^26^. The X and Y dimensions of the volume (630 nm) depend on the magnification, while Z (150-250 nm) approximates the physical thickness of the sample. The thicker the sample, the lower the SNR. **c**. A slice through the reconstructed volume before and after denoising. The thickness of the slice is equal to 5 Å (Ångstroms = 10^-10^ meters). One example for each of the 5 species that are the targets of the challenge is indicated by an arrow. The colors correspond to the colored structures in panel d. **d**. 3D reconstructed volumes of the protein complexes that are the targets of the challenge. These volumes/maps are from published data (refer to Table 1, also refer to Extended Data Fig. 5 for the maps reconstructed from the phantom data). In a common cryoET workflow, for each species thousands of copies of that protein are annotated, extracted, and further processed to produce a high-resolution map. These high-resolution maps reveal how the proteins are organized in space and how they carry out their functions.

**Table 1.** Purified proteins (from different sources) present in the phantom dataset. All proteins except for Human Serum Albumin (HSA) and Beta-amylase are targets in the machine learning challenge. The listed EMD entries are for the maps used in Figures 1 and 2, and some initial picking with PyTom template matching.

| Sample | MW | Symmetry/copies | Average counts per tomogram | ID |
| --- | --- | --- | --- | --- |
| Ribosome (80S; human) | 4.3 Md | Monomer<br>1 x | 80 | EMD-3883 |
| VLP (PP7) | 3.4 Md | Icosahedral<br>60 x | 6 | EMD-41917 |
| Thyroglobulin (bovine) | 660 Kd | Homodimer<br>4 x | 13 | EMD-24181 |
| Beta-galactosidase | 540 Kd | D2<br>4 x | 6 | EMD-0153 |
| Apoferitin | 450 Kd | Octahedral<br>24 x | 47 | EMD-41923 |
| Beta-amylase (Sweet potato) | 268 Kd | D2<br>4 x | 5 | EMD-30405 |
| Albumin (Human) | 66 Kd | Monomer<br>1 x | NA | EMD-43090 |

However, there are several challenges to obtaining high resolution maps of protein complexes with cryoET. First, unlike CT and MRI, in cryoET the range of projection views is restricted to less than +/- 60° (Extended Data Fig. 1a) which results in artifacts in the reconstructed volumes, commonly known as the missing wedge artifact^10^. Second, because high energy electrons are damaging to biological samples, the number of electrons that can be used is severely limited and thus the resulting images and volumes have very low signal to noise ratios (SNRs) (Fig. 1c). As a result, we need to identify and average together thousands of identical entities of different orientations from many 3D volumes in order to produce a high resolution map of a protein complex of interest^1^ (Fig. 1d). The process of identifying these individual entities of interest in the noisy 3D volumes is referred to as annotating, picking, or labeling.

Annotating cryoET tomograms remains a significant bottleneck due to the low SNRs of the projections and volumes, the structural complexity of cellular samples, the diversity and heterogeneity of the molecules of interest, and the large numbers of molecules that are required for high resolution maps. In most cases, annotation is the most time-intensive and laborious part of cryoET data processing since it often relies heavily on manual input. Comprehensive labeling is critical both to obtaining high-resolution structures of individual protein complexes and understanding the organization of large-scale ultra-structures, so new methods to annotate cellular tomograms at scale are urgently needed.

Machine learning (ML) algorithms are well-suited to overcome this annotation bottleneck^11–15^. At the time of writing, 15,732 tomograms are publicly available through the recently launched cryoET Data Portal^16^ (cryoetdataportal.czscience.com). While machine learning has been leveraged to provide membrane segmentations for all datasets in the portal, labeling particles is a far more difficult task due to their diversity, lower contrast, and crowding. As a result only 5% of tomograms in the portal have molecular annotations. Further, labeling strategies that can be readily adapted to account for data characteristics that vary with acquisition parameters and better capture the intrinsic heterogeneity of molecules would be advantageous compared to traditional approaches. ML methods to date are effective for the specific cases they were developed on, but typically do not generalize sufficiently to meet the diverse needs of the cryoET community.

Previous machine learning challenges, both in cryoEM and cryoET, have been valuable in benchmarking several algorithms using simulated datasets^17–19^ or limited real-world datasets^20^. Here, for the first time, we have organized a machine learning challenge (https://cryoetdataportal.czscience.com/competition) based on a real-world diverse sample and a large cryoET dataset to spur innovation in this domain. Contestants will be tasked with developing ML algorithms that can robustly perform multi-class particle labeling on hundreds of unseen tomograms after being trained on a limited set of annotated tomograms from the same dataset, as well as any other data — either synthetic or experimental — of the competitors’ choice. To encourage generalizability, we have chosen five target particles that have diverse shapes and collectively span nearly an order of magnitude in molecular weight (Fig. 1d). Four target particles, Virus-like Particles^21^ (VLPs), Thyroglobulin (THG)^22^, Beta-galactosidase^23^, and Apoferritin^24^, were mixed with cellular lysate, which naturally contains the fifth particle, 80S ribosomes^25^ in abundance (Fig. 2a). The lysate also naturally includes various non-target particles, such as nucleosomes, filaments, proteasomes, and membrane-bound proteins, along with structural elements like membranes, which frequently confuse picking algorithms (Extended Data Fig. 2). We generated reference annotations through an elaborate and rigorous workflow, as described below, that drove the development of several new tools to aid particle picking but also underscored the need for a more streamlined and fully automated solution to tomogram annotation. These “ground truth” labels will be used to score participants’ results and will be released on the CryoET Data Portal as a resource to benchmark future algorithm development after the contest ends. We anticipate that this challenge will deliver novel algorithms to streamline tomogram annotation, provide a benchmark dataset and a standardized pipeline to guide the continuous improvement of ML models, and engage the ML and cryoET communities to jointly tackle challenges facing the field.

**Figure 2.**
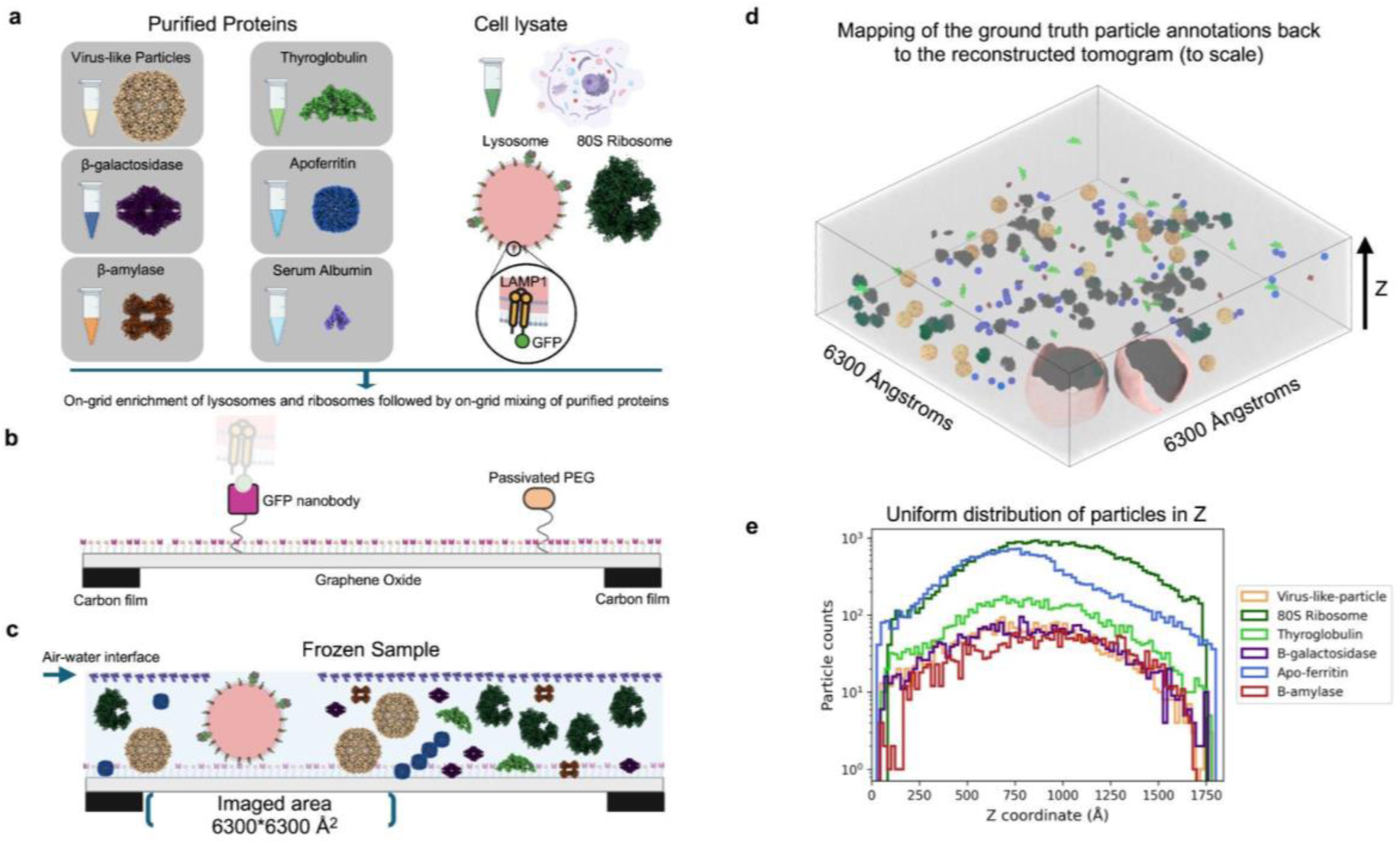
The composition of the phantom sample. **a**. Two sources were used in the preparation of the phantom sample: purified proteins and cell lysate. From the cell lysate, lysosomes were captured by affinity interactions between LAMP1-GFP and GFP nanobody. Ribosomes co-purified because of their high abundance. **b**. Cross-section (1.2-µm width) of the functionalized EM grid surface prior to sample application. The carbon film (20-nm height) serves as a support for, from bottom to top: graphene oxide (1-atom thickness), combination of passivated PEG and PEG-maleimide-GFP-nanobody. **c**. A schematic cross section through the sample on the grid; note proteins are not drawn to scale. The proteins are frozen in time in glass-like ice (ice without crystals). Tilt series were collected with a field of view of 6300 Å x 6300 Å in regions excluding the carbon film. **d**. A 3D view of a typical phantom tomogram. Annotations drawn to scale. All the annotated particles (except for HSA) are placed in their true positions in a tomogram with the boundaries of the 3D volume outlined. The membranes are rendered as surfaces. **e**. The distribution of all the annotated species in Z shows an almost uniform distribution across the thickness of the sample indicating the advantage of using the functionalized grids to avoid all proteins adhering to the air-water interface.

### Creating a “phantom” sample

Prior to setting up the machine learning challenge, we carried out a survey of interested parties, which indicated that there was limited enthusiasm for basing a challenge around synthetic datasets nor for the several well annotated existing ribosome datasets. Inspired by the work of Ishemgulova et al^27^, we opted to create a sample from a mixture of cell lysate and known proteins. We refer to this created sample as a “phantom”, based on nomenclature used in the biomedical research community^28^ to denote objects used as stand-ins for tissues to evaluate systems and methods for imaging. The primary reason for using this phantom rather than cellular material is the inherent complexity and time-consuming nature of preparing and annotating cell-based samples. Working with intact cells introduces a multitude of variables and intricate structures that are difficult to label exhaustively or accurately. This makes it exceptionally challenging to create “ground truth” datasets suitable for training machine learning models and scoring contestants’ results. Therefore, we developed this phantom to mimic the cellular environment to the extent that it provides a relatively large set of annotations for diverse protein structures. For users who are interested in tuning machine learning algorithms to the crowded *in situ* datasets, the cryoET data portal (cryoetdataportal.czscience.com), discussed below, provides multiple annotated datasets of protein complexes *in situ*.

Our phantom sample comprises several key components designed to simulate a reduced level of the diversity and complexity of a cellular tomogram (Fig. 2). Firstly, we utilized HEK293T cell lysate^29,30^ that serves as a foundational component, offering a realistic backdrop of cellular material, including common elements, such as ribosomes and membranes (Fig. 2a). To further diversify the sample, we mixed in five commercially available proteins, apoferritin, thyroglobulin (THG), Beta-galactosidase, Beta-amylase, and Human Serum Albumin (HSA), as well as virus-like particles (VLPs) (see Acknowledgements). These components collectively provide a range of molecular weights, shapes, and therefore complexity (Table 1) for testing and developing machine learning algorithms for protein complex annotation.

To produce the phantom sample, we used functionalized electron microscopy grids^31,32^ optimized for cell lysates. The surface of the grid was covered with a layer of graphene oxide (Fig. 2b) and then functionalized with GFP-nanobodies that bind to a GFP-Tag attached to the LAMP1 C-terminus found on lysosome surfaces in HEK293T cells (Extended Data Fig. 1b). This design selectively enriches lysosomes on the grids while other larger organelles such as mitochondria are washed away. However, because of the “gentle” washing step, abundant cellular protein complexes such as ribosomes and nucleosomes remain on the grid. The binding of lysosomes onto the graphene oxide layer creates a spacer effect (Fig. 2c, 2d) that facilitates a consistent sample thickness (∼150-250 nm) similar to that typically seen in cellular samples. Sample thickness is a critical factor in cryoET as samples that are too thick have reduced contrast, while samples that are too thin provide very limited cellular volumes. Using our sample preparation method we were able to achieve two features that are otherwise improbable: 1) multiple layers of particles stacked on top of each other (Fig. 2d, 2e, Extended Data Fig. 3) similar to *in situ* samples, 2) exclusion of most of the proteins of interest from the air-water interface (Fig. 2d, Extended Data Fig. 4) which can cause protein denaturation or induce preferred orientations. In these phantom samples, we observed that the air-water interface is consistently populated by very small protein densities (Fig. 2c, Extended Data Fig. 4). We hypothesize these densities to mostly belong to the added HSA and Beta-amylase, as well as small proteins in the lysate. Among the larger proteins, a small subset of ribosomes partially makes contact with the air-water interface. Crucially, preventing preferential orientation allows for a more accurate representation of molecules (Extended Data Fig. 5), which is essential for training robust machine learning models.

Our phantom sample captures many of the essential elements of cellular cryoET data, including inherent noise and contrast, a range of shapes and sizes of protein complexes, and the challenges associated with distinguishing species with similar-looking shapes. For example, this similarity arises when the small dimension of a larger protein has similar features to the large dimension of a smaller protein, so that an algorithm interprets the similar views as belonging to the same class of proteins (compare the 2D class views of THG, Beta-galactosidase, and Beta-amylase from Extended Data Fig. 5). The phantom sample also reflects the heterogeneity seen in ribosomes from cells in various states and includes membranes and membrane-bound proteins, although these are not the primary focus of the current challenge. We also observe low copy numbers of additional cellular structures, such as nucleosomes, proteasomes, and lumenal proteins which are scattered sparsely through the tomograms, increasing the complexity of the sample (Extended Data Fig. 2).

It is important to note however, that our phantom sample does not capture the crowdedness of real cellular environments. While it includes a variety of components, and mimics many aspects of cellular complexity, the density and spatial constraints typical within a living cell are not replicated. While this limitation is acknowledged, the phantom sample nevertheless serves as an initial well annotated dataset for developing, training, and advancing specialized machine learning models for cryoET data analysis, with the anticipation that this will lead to algorithms capable of labeling protein complexes in realistic cellular environments.

### Establishing “ground truth” labels

Ground truth labels, here, are defined as the *x,y,z* coordinates of the center of each particle in its tomogram with the origin of the coordinate system being the corner of the tomogram. Generating “ground truth” labels required multiple months of work by a large team of people and spurred the development of several new methods for particle picking, visualization, and curation (Fig. 3, Extended Data Fig. 6). This effort included manual annotation, the use of template matching and machine learning algorithms, and the careful curation of the selected particles using a variety of approaches. Below we will first briefly outline the in-house tools that were developed as part of these efforts; these will be described in more depth in forthcoming publications (Table S1). We have already repurposed several of these tools for other projects at our institute so we anticipate these new methods will prove valuable beyond this challenge.

**Figure 3.**
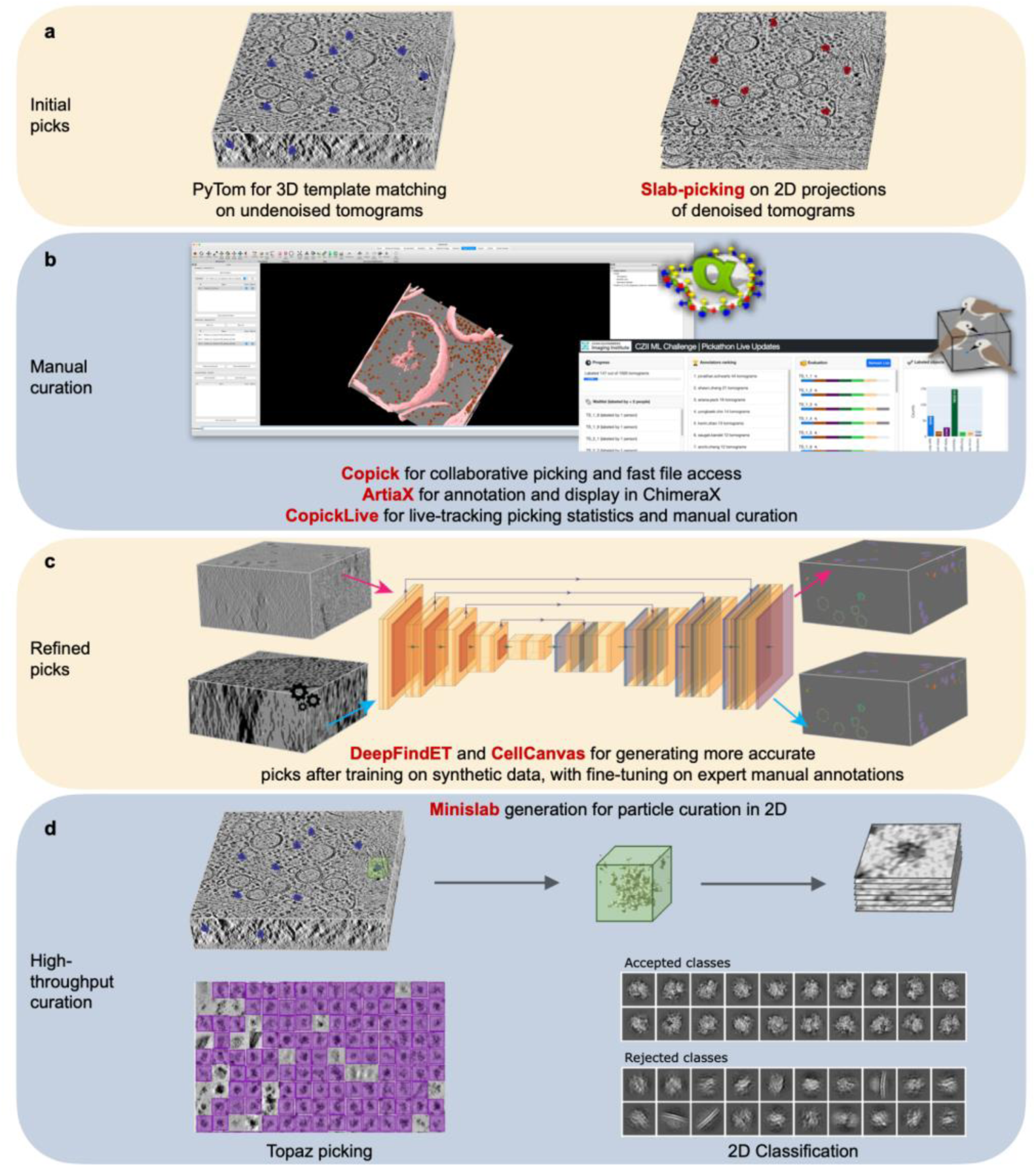
Multi-step workflow developed to generate ground truth labels. **a**. Particle picks were initially generated either by PyTom^33,34^ 3D template matching algorithm or 2D slab-picking from 300 Å thick projections through denoised tomograms. **b**. These picks were manually curated in ChimeraX^35^, using ArtiaX^36^ for annotation, copick for rapid file access, and copicklive to track the progress of curation. **c**. Two ML pipelines, DeepFindET and CellCanvas, were trained on synthetic tomograms and fine-tuned on a selection of the curated picks from expert annotators to generate a more refined and complete set of picks. **d**. Final rounds of curation were performed by applying Topaz, 2D classification, and multi-class *ab initio* reconstruction to per-particle 2D projections (“minislabs”) generated from denoised tomograms to filter out contamination and non-target particles. New tools developed as part of this curation effort are indicated in red.

*DenoisET* implements the *Noise2Noise* algorithm^37^ for denoising cryoET tomograms. This ML algorithm learns to denoise imaging data from training on paired noisy measurements and is thus suitable for methods like cryoET in which the clean signal is difficult to realistically simulate and cannot be measured. However, because cryoET data are collected as a series of frames of the same underlying field-of-view but with different realizations of the stochastic noise process, training data can be readily generated by reconstructing paired tomograms after splitting the acquired frames into half-sets. Our implementation relies on a similar U-Net architecture as used in Topaz-Denoise^38^ and leverages the contrast transfer function-deconvolved tomograms produced by AreTomo3^26^ to improve contrast enhancement. The increased SNR provided by denoising proved critical for our manual annotation efforts and benefited our machine learning labeling workflows as well (Fig. 1C).

*Copick* is a storage-agnostic and server-less platform designed specifically for cryoET datasets. This package permits efficient access of tomograms and annotations both programmatically, to assist algorithm development, and visually in ChimeraX^35^ and napari^39^ for inspection and manual labeling. Specifically, tomograms are stored in the multi-scale OME-Zarr format^40^ to enable rapid and parallel data loading from any filesystem at different resolutions. Annotations are associated with a particle class and other unique identifiers and can be overlaid in an editable format on static copies of reference labels. Collectively, these design choices enabled a large team to manually curate our initial picks in parallel and simplified methods development by providing a unifying framework for data access and storage. We leveraged this framework to easily transfer annotations among all of the in-house tools described here.

*Slab-picking* was developed as an alternative to 3D template matching to generate initial candidate picks (Fig. 3A). For this approach, tomograms were divided along the z-axis into slabs of uniform thickness and projected along that axis to increase the SNR. These projections were then uploaded into CryoSPARC^41^ as mock micrographs. Conventional 2D approaches for particle picking such as blob-picking and template matching were applied to locate candidate particles in each micrograph, and 2D classification was performed to remove false positives by manually deselecting classes judged to be incorrect. The locations were then mapped back to their corresponding tomograms, and the z-height of each particle was refined based on local intensity statistics. Due to time constraints, we were limited to generating candidate picks for all particle types from non-overlapping 300 Å thick slabs. However, we expect that using a sliding window approach to retain particles positioned at the slab boundaries and adjusting the slab height to match the particle size would yield more hits.

*ArtiaX*^36^ is a ChimeraX^35^ plug-in that provides a toolbox for visualizing, selecting, and editing particle picks in tomograms (Fig. 3B). This package was extended in several ways to facilitate and accelerate large-scale manual curation efforts. First, the package was rendered interoperable with the copick framework to enable annotators to rapidly render tomograms at different locations and at distinct voxel spacings, quickly switch between tomograms of interest, and independently curate the same set of candidate picks in parallel. Second, shortcut keys were added to scan candidate picks by recentering the field-of-view on each particle and rapidly delete false positives. Finally, a new feature provides orthoslice views through the tomogram to aid identification and disambiguate particles that are indistinguishable in projection.

*Copicklive* provides an interactive web viewer to track picking statistics and tomogram curation in real-time (Fig. 3B). These features proved useful for maximizing annotation coverage across the full set of tomograms and reducing duplicate efforts during a week-long manual picking marathon that included 42 individual participants who collectively annotated 147 tomograms and over 29,000 particle annotations. In addition, the interface provides tools to facilitate particle rejection and class reassignment based on 2D projections of subvolumes centered around candidate picks. Because copick is the foundation of this software package, the results of manual curation are automatically saved in a standardized format that can be easily accessed by other software.

*DeepFindET* is an adaptation of the *DeepFinder* package^12^, a CNN-based algorithm to simultaneously label multiple particle types in cellular cryoET data. While initial results from using this package out-of-the-box were promising, the following extensions significantly improved performance on our phantom dataset. First, the model was switched from a U-Net to Residual U-Net architecture for more stable training. Second, additional geometric-based augmentations were added to diversify the training data. Third, the original semi-supervised clustering approach was replaced with a size-based selection scheme to reduce bias and increase confidence in the particle labels during inference. The model was trained on synthetic data containing a mixture of target particles and a limited set of phantom tomograms labeled by expert annotators (Fig. 3C).

*CellCanvas* is a flexible tool for building geometric models of cellular architecture, with an associated napari plug-in to enable interactive painting-based segmentation and model refinement (Fig. 3C). CellCanvas’ segmentation capabilities were used to generate an intentionally over-picked set of candidate particles to minimize false negatives. The first step of this pipeline generated a voxel-wise embedding for each tomogram in the phantom dataset using a Swin UNETR model, which was pre-trained on medical imaging data^42^ and fine-tuned on four synthetic tomograms generated using PolNet^43^. Clusters in embedding space were transformed into segmentation masks by interactively training an XGBoost classifier on manual annotations. These segmentation masks were in turn converted to particle picks. While we found the combination of pre-computed embeddings and quick-to-train interactive models highly effective for labeling the phantom data, CellCanvas was intentionally built with a plug-and-play design that enables replacing the embedding model and classifier with other models for increased flexibility.

*Minislab curation* was developed to facilitate particle curation which is the filtering of picked particles. In this approach, subvolumes centered on individual particles were extracted from the tomograms, and the intensity was integrated along the z-axis to generate per-particle 2D projections (Fig. 3D). These “minislabs” were then tiled to generate mock micrographs for further processing in CryoSPARC^41^. The particles were cleaned up and curated by various combinations of performing manual picking, 2D classification, running *ab initio* reconstruction with multiple classes, and applying Topaz^14^, a CNN-based particle picker that was retrained for each particle class. The selected particle projections were mapped back to their positions in the tomograms. While minislab curation cannot distinguish between particles that appear similar in projection, curation of 2D projections was more efficient and benefited from the increased signal-to-noise compared to 3D subvolumes.

The tools described above, and other published software packages were stitched together into several workflows to generate “ground truth” labels (Fig. 3, Extended Data Fig. 6). Initial picks were generated either using PyTom^33,34^ 3D template matching (VLP, ribosome) on undenoised tomograms or 2D slab-picking (apoferritin, Beta-galactosidase, thyroglobulin) on denoised tomograms. Preliminary picks for a subset of tomograms were manually curated in ChimeraX^35^, leveraging new features in ArtiaX^36^ for facile annotation and copick for rapid file access and exchange. These picks were curated by expert annotators for the next step of model training. To generate a more refined set of picks across the full phantom dataset with less false negatives or missed particles, DeepFindET and the CellCanvas classifier were trained on a combination of synthetic tomograms generated by PolNet^43^ and real data curated by the expert annotators. The PolNet training set contained the five particles of interest in addition to membranes and a sixth particle, Beta-amylase. Minislabs for the predicted picks were extracted from denoised tomograms for final rounds of curation. For VLP and apoferritin, curation was achieved by applying a Topaz model trained separately for each particle. For thyroglobulin and Beta-galactosidase, 2D classification was performed and the retained picks from both CellCanvas and DeepFindET were merged into a single set. After removing duplicates, the picks underwent a round of manual curation followed by multi-class *ab initio* reconstruction to reject any remaining false positives. For the ribosome, both Topaz and iterative rounds of 2D classification were used to eliminate contamination. Given the intrinsic heterogeneity of ribosomes, we performed final rounds of 2D classification and multi-class *ab initio* reconstruction to provide a confidence score for each retained particle. In the case of particles curated by Topaz, any true positives rejected by the model were rescued through manual curation to minimize false negatives. This long, complex, and convoluted approach required to generate accurate annotations highlights the need for more streamlined algorithms that generalize well across datasets and diverse target particles and thus the urgent need for this ML challenge.

While we refer to our final curated set as “ground truth”, we acknowledge the following caveats. Based on our rigorous curation efforts, we are confident that the less challenging particle labels contain few false positives but are less confident about the more challenging particles. We also acknowledge that our ground truth labels very likely missed some true positives. These caveats are due to the fact that even a well-trained expert finds it exceptionally difficult to be certain in labeling the smaller particles and it would take years for experts to fully annotate this large dataset. We also note that we opted for a more inclusive set of particles than would be included in the typical downstream processing task of subtomogram averaging. Thus, our final labeled particles are conformationally heterogeneous and include partial particles at the tomogram boundaries if their identities seem clear. In the case of the VLPs, we retained not only the abundant 29 nm diameter icosahedral form but also other spherical particles and tubular species. Our intention in making these choices was to focus the challenge on the specific task of labeling without constraining the labels to the most homogeneous subset of particles in the data. The final ground truth set includes thousands of picks for each of the THG, Beta-galactosidase, and VLP particles, and tens of thousands of picks for each of the 80S ribosome and apoferritin particles (Table 1, Extended Data Fig. 5).

### The Machine Learning Challenge

The cryoET field has benefited from an increase in machine learning contributions, but particle picking, a problem that is well-suited for machine learning methods, remains a major open problem. We are hosting an ML challenge (https://cryoetdataportal.czscience.com/competition) to push the limits of particle picking across particle sizes and draw attention from the broader ML community. We deliberately selected a small training subset of the full dataset to recapitulate the common situation of a cryoET researcher who can only afford enough effort to annotate a handful of tomograms, yet needs annotations for hundreds or thousands of tomograms for high-resolution subtomogram averaging. We have designed evaluation metrics that span a range of particle sizes, incentivizing models that can perform well for small particles. We note that due to the challenges of annotating particles, even by well-trained human experts, there is the possibility that a fraction of the particles have not been labeled in the ground truth. These false negatives will be explored after the competition is over. We will use a variety of 2D and 3D classification methods to assess the particles provided by the top 10 submissions to determine if they include a larger fraction of true positives than the ground truth we provide. This analysis will be shared with the community and published as part of a larger assessment of the ML challenge results, the winning algorithms, and the lessons learned.

#### Data description

The 3D volumes provided to the participants on Kaggle-for training and testing-are denoised tomograms (Table 2). The three other tomogram types (weighted back projection, CTF-corrected, Isonet-corrected) are available on Kaggle only for training and not testing. All the tomogram types are available to the participants on the cryoET data portal (as well as raw data such as frames and tilt series). The ground truth labels for these tomograms are *x,y,z* coordinates of the center of each labeled particle. The origin for this coordinate system is the lower bottom left corner of each tomogram (which would be increasing positively along the x, y, and z axes of the 3D image array). These ground truth labels belong to 6 distinct classes which are evaluated individually. Therefore, participants are required to submit their labels in the same x,y,z coordinate system per class.

**Table 2.** Different tomogram types available in the challenge. All of these tomograms have the same tilt series alignment, therefore, the particle coordinates are the same across all types.

| Tomogram Type | Reconstruction workflow | Description | Contrast |
| --- | --- | --- | --- |
| Weighted Back Projection | AreTomo3 | Real space WBP | Low |
| CTF-Deconvolved | AreTomo3 | Local CTF correction of tilt series+<br>real space WBP | Medium |
| Denoised | AreTomo3<br>+DenoisET | Local CTF correction of tilt series+<br>real space WBP+tomogram denoising | High |
| Isonet-Corrected | AreTomo3<br>+DenoisET<br>+Isonet | Local CTF correction of tilt series+<br>real space WBP+tomogram denoising+missing<br>wedge estimation and correction | High |

#### Evaluation metric

The evaluation metric for the cryoET particle picking challenge uses a distance threshold-based point matching approach, where matches are determined based on a multiple of the particle radius. The metric employs microaveraging, averaging matches across all picks rather than per tomogram. To mitigate the penalty for false positives, an F-score with a beta value of 4 is utilized, emphasizing recall and reducing the penalty for false positives given the annotation uncertainty for small particles (e.g. recall is 16 times more important than precision). Additionally, the metric incorporates weighted scoring for different particle groups, assigning higher importance to hard-to-pick particles (Fig. 4). A weight of 1 is assigned to the “easy” to pick particles: apoferritin, ribosome, and virus-like particles (VLPs), while a weight of 2 is assigned to the more difficult particles to pick: thyroglobulin (THG) and Beta-galactosidase. A weight of 0 is assigned to Beta-amylase, because it’s not part of the challenge, although annotations are provided for training purposes.

**Figure 4.**
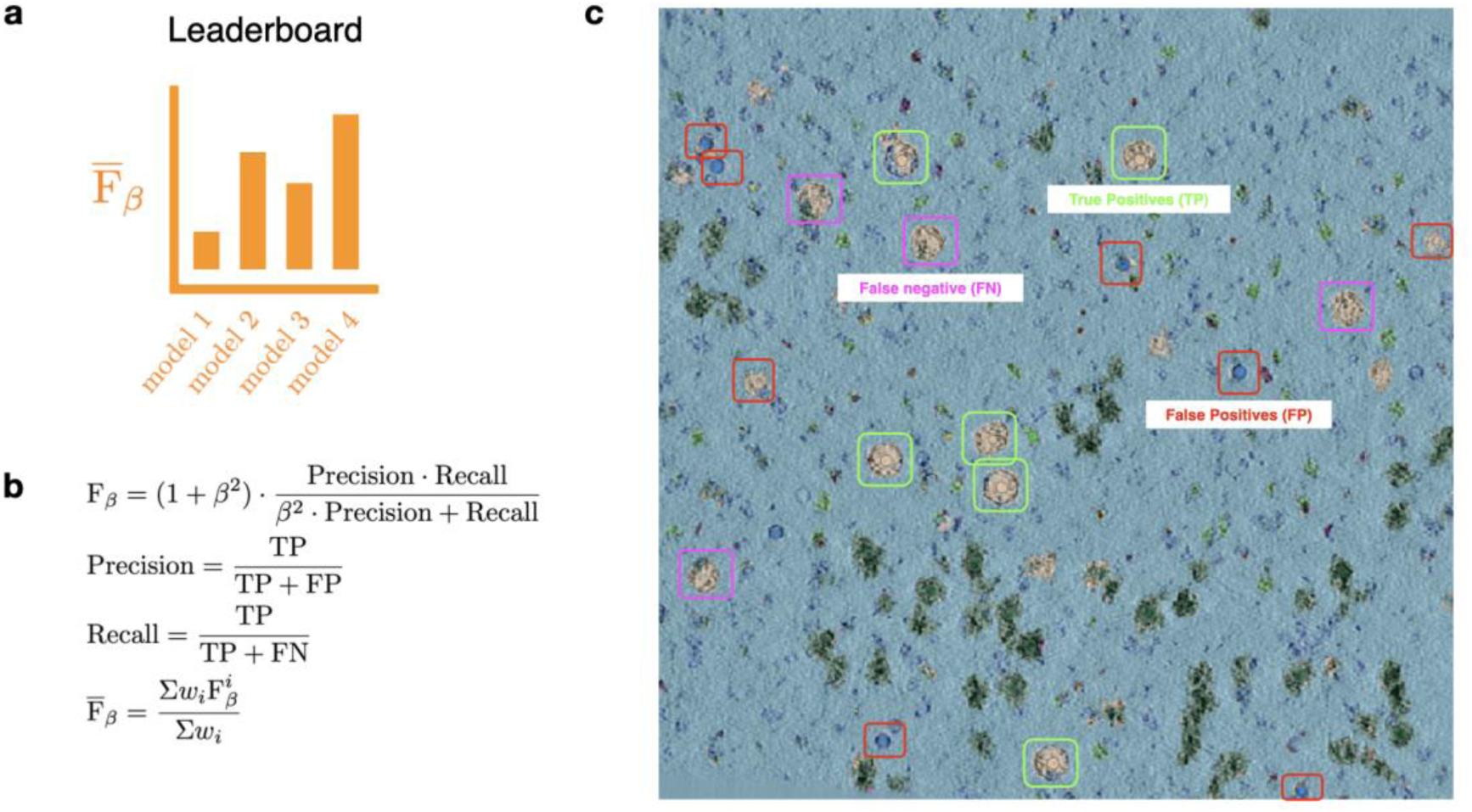
Evaluation of models’ picking performance. **a**. The leaderboard ranking is determined by **b**. the weighted average F_beta score, where the F_beta score is first calculated for individual protein species and then combined across all five species using weighted aggregation. **c**. A tomogram slice annotated in CellCanvas demonstrates model detection outcomes. True positives, where the model correctly identifies VLPs, are marked with green squares. Missed VLPs, or false negatives, are indicated by magenta boxes, while structures mistakenly identified as VLPs (false positives) are highlighted in red.

**Figure 5.**
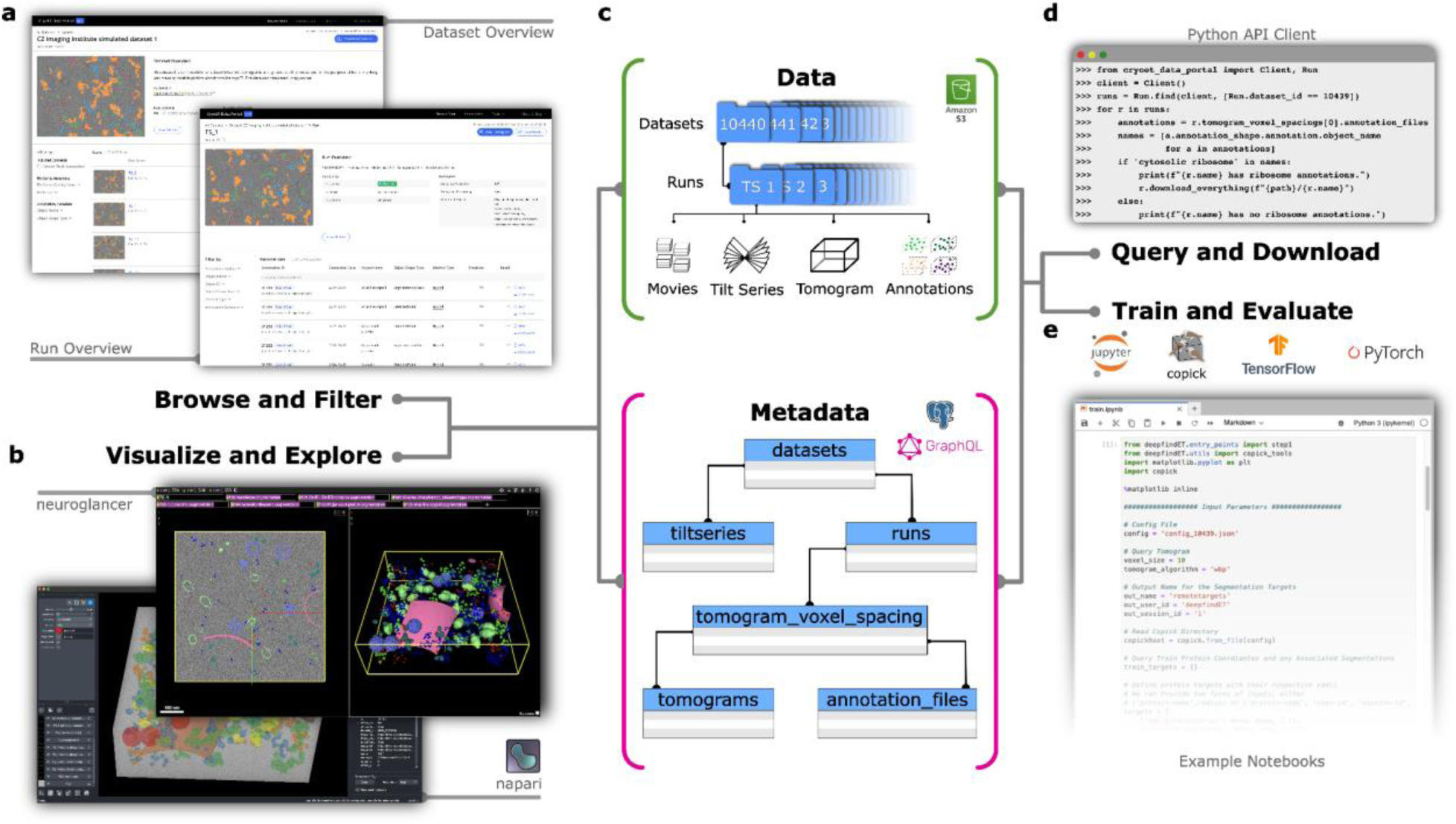
The cryoET data portal and related infrastructure. **a**. The cryoET data portal web interface allows challenge participants to explore the experimental and simulated Challenge datasets (dataset IDs 10440 and 10441), included runs, annotation and their metadata. **b**. Tomograms and annotations can be visualized directly inside the web browser using Neuroglancer^48^, or in the desktop viewer napari^39^. **c**. All related image data and metadata are provided from a public AWS S3 bucket and associated metadata can be queried using a publicly exposed GraphQL API**. d**. Both resources can be accessed programmatically using the cryoET data portal python API client, which also provides convenient methods for data download. **e**. All cryoET data portal resources (including other datasets) can be accessed using copick and the provided example notebooks, which feature reference implementations in TensorFlow^47^ and PyTorch^46^.

#### Dataset split

We curated a selection of 492 good-quality tomograms (Extended Data Fig. 7) from the phantom dataset and divided them into three subsets: training, testing, and validation. To ensure that these subsets had similar distributions across all particle species, we employed the Kolmogorov-Smirnov test to assess the similarity between distributions. The 492 tomograms were then grouped into 16 bins based on this analysis, and tomograms randomly sampled from these bins were used to construct the three subsets. We experimented with various data split ratios and observed that DeepFindET was able to perform well with smaller training sets, which is the most common situation in CryoET. Consequently, we opted to submit only seven tomograms to be used for training, 25% of the remaining tomograms are used for public evaluation and 75% are used for private evaluation and final scoring of the submitted models

#### Establishing metrics parameters with synthetic datasets

To ensure that the F-beta scores effectively distinguish performance differences among models (especially models that will perform better than DeepFindET), we determined the metric’s beta value using a set of results with known ranking quality. This set was generated by mixing varying percentages (in 10% increments) of picks from a fine-tuned DeepFindET model and ground truth data. By comparing their F-beta scores, we observed that datasets containing 80% or more ground-truth results yielded mock datasets that score better than DeepFindET. Based on this insight, we added an additional 20 datasets ranging from 80% to 100% (in 2% increments) ground truth with the remaining picks coming from DeepFindET and used them to compare the impact of different beta values. For this comparison we aimed to find the smallest beta value that ensured a consistent top-5 ranking. Through this analysis, we determined that setting beta=4 produced a more consistent ranking.

#### Example notebooks

To reduce the onboarding time for competitors, an extensive set of example notebooks is being provided. These include: DeepFindET, TomoTwin, and 3D U-Net models. For each model type (except TomoTwin which uses a pretrained model), training and inference notebooks are provided. These notebooks also leverage the copick library for handling cryoET datasets, for which we provide PyTorch Datasets and utility functions to simplify the creation of data loaders, metadata tracking, and model performance analysis.

### Enhancing training infrastructure and data using the cryoET data portal

All training data described here (7 tomograms) as well as additional published training data are available on the CZ CryoET Data Portal (CZCDP^16^, cryoetdataportal.czscience.com) under dataset IDs 10440 (experimental data) and 10441 (PolNet^43^-simulations). The CZ cryoET Data Portal is an open platform specifically designed to facilitate algorithm development, including by researchers outside the cryoET field. It expands on the data archiving and sharing efforts made by the Electron Microscopy Public Image Archive (EMPIAR)^44^, and the Electron Microscopy Data Bank^45^, by providing a large collection of curated and standardized datasets and annotations under public domain (Creative Commons 0) license, as well as a growing set of open source tools geared towards machine learning development and cryoET visualization.

Independent of domain expertise, ML Challenge participants can familiarize themselves with the training data, as well as other available cryoET datasets, by browsing the Portal’s web interface and visualizing the provided ground truth annotations in their web browser using the integrated Neuroglancer-based viewer. Outside of the browser, datasets can be queried using a GraphQL-API, accessed using a python-based API client and visualized using a plugin to the ND-image viewer napari^39^. In contrast to existing archives, all datasets are accessible with a consistent layout and metadata schema in public cloud storage, and directly link raw data, tilt series, tomograms and annotations. All processed image data are stored in the cloud-ready next generation file format OME-Zarr^40^, allowing participants to train particle picking networks on datasets exceeding available local storage by streaming tomograms whole or in part, and removing dependencies on domain-specific image formats (such as MRC). PyTorch^46^ and Tensorflow^47^ infrastructure relying on these features is available to participants through the direct integration of CZCDP datasets with the copick API.

We expect that the considerable volume (15,732 tomograms) and diversity (44 species) of the data already hosted on the CryoET Data Portal will lend itself to the development of robust ML algorithms that generalize well across samples. In addition, the growing collection of datasets should serve as further motivation to ML Challenge participants, as any tools and algorithms relying on CZCDP infrastructure and/or copick will be immediately applicable to the entire current collection, as well as any future data deposited to the portal, with little or no need for modification.

Taking into account the available datasets and infrastructure, we envision direct ways through which the cryoET Data Portal could be leveraged to improve model training for this challenge. For example, the synthetic data used to train CellCanvas and DeepFindET have been made available (portal dataset ID: 10441), and can serve as convenient test sets for models and infrastructure. In addition, although these synthetic tomograms lack the complexity of experimental data, we observed that training on them significantly improved prediction accuracy across all particle classes for both of the models we used. The deposited annotations may also prove a valuable resource for model refinement. In particular, the 200,843 instances of non-mitochondrial ribosome labels across 447 non-phantom tomograms could supplement the limited training tomograms from the phantom dataset. Although this is the only target particle that has been annotated in other Portal datasets, including molecular annotations for non-target particles could potentially improve a model’s ability to distinguish among different molecular species. In a similar vein, the membrane segmentations available for all Portal datasets could be used as negative training examples to prevent algorithms from mistaking these high contrast features for target particles. We anticipate that the specific ways in which contestants incorporate data and annotations from the CryoET Data Portal into their workflows will be informative for future algorithm development.

A final distinguishing feature of the CZ cryoET data portal is the evolving nature of its corpus. Annotations generated as part of the Challenge will be added to the cryoET data portal with attribution to the authors and direct links to their tools and methods. This provides a simple path for participants to present their results interactively to a broad audience and allows for easy reproducibility and comparison of any methods developed as part of the Challenge.

## Discussion

Challenges like the one described here have catalyzed the development of state-of-the-art methodologies in structural biology. A particularly instructive example is the biennial Critical Assessment of protein Structure Prediction (CASP) competition that benchmarks protein structure modeling algorithms against experimentally-determined structures^49^. Compared to the incremental progress achieved during CASP’s first decades, the introduction of DeepMind’s AlphaFold in 2018 led to a major breakthrough in prediction accuracy^50^. This leap in performance was so profound that CASP has since shifted its focus to more challenging protein modeling problems, while AlphaFold has increasingly been integrated into other structural biology workflows^51,52^. This advance critically depended not only on innovations in deep learning research but also on the public availability of a rich corpus of experimental results^50,53^.

Until recently, the conditions to hold a Machine Learning Challenge had not been met in the field of cryoET. However, several developments have converged to make such a competition not only possible but potentially game-changing for the field. As with protein structure modeling, a wealth of experimental data is now available across multiple public data banks. EMPIAR^44^ and EMDB^45^ have led the charge to make cryoET data openly accessible. Expanding on these efforts, we launched the CryoET Data Portal to prioritize data standardization, tomogram annotation and segmentation, including by non-cryoET practitioners so that domain-specific knowledge is not a barrier to entry. The recent acceleration of data acquisition and processing is poised to vastly expand the quantity of public data and it has also enabled us to generate a benchmark dataset that includes hundreds of high-quality tomograms. Importantly, this phantom dataset contains artifacts characteristic of experimental tomograms and is large enough to robustly score annotation algorithms in blind predictions against withheld data. As the existing labeling approaches tried in this project did not work out-of-the-box on our phantom tomograms, rigorously annotating the full dataset required months of effort and stitching together in-house tools and existing software into a convoluted workflow that underscored the need for a more streamlined solution. The result of this effort is high-quality annotations across nearly five hundred experimental tomograms for six distinct particle classes whose range of shapes and sizes will identify deep learning algorithms most likely to generalize to the diverse targets of interest to the cellular biology community.

We expect that our Machine Learning Challenge will serve as the foundation for a series of contests designed to spur innovation in cryoET methods development, including from experts outside the field, and provide a standardized platform to critically evaluate state-of-the-art annotation algorithms. The results from these contests will in turn inform the design of future benchmark datasets that best showcase the capabilities and identify the limitations of current methods, offering the field a valuable resource to continually track its progress. Similar to CASP, advances will enable us to expand the scope to more difficult challenges such as annotation in crowded cellular environments and labeling molecular features along high-contrast membranes. We hope that these challenges over time will deliver transformative ML methods that solve the annotation bottleneck in cryoET and can be readily scaled across all available data to deepen our insights into how molecules are structured and organized in their native context of the cell.

## Methods

### Sample preparation

The phantom dataset was generated from a combination of cell lysates, commercially available purified protein, and purified protein from collaborators. The cell lysates were from a HEK293T cell line generated at Chan Zuckerberg BioHub, San Francisco by Manuel Leonneti’s group with a knock-in GFP-Tag on the C-terminus of the lysosomal house-keeping protein LAMP1. The homogenization protocol was developed at our collaborator’s lab^30^ and adapted by us for cryoET sample preparation. Briefly, the cells were lysed using a hypotonic homogenization buffer (25mM Tris*HCl pH 7.5, 50mM sucrose, 0.2mM EGTA, 0.5mM MgCl_2_) and shearing forces generated using a 23G syringe. To protect the organelle membranes the lysate was immediately mixed with a sucrose buffer (2.5M sucrose, 0.2mM EGTA, 0.5mM MgCl_2_) to re-equilibrate the homogenates to an isotonic osmolarity. The nuclear fraction was separated by centrifuging the lysate at 1000 x g for 10 minutes. The lysosomes with the LAMP1-GFP tag from this lysate were then purified on-grid using functionalized electron microscopy grids^31,32^. This technology was adapted for organelles at the CZ Imaging Institute and University of California, San Francisco in a joint collaboration with David Agard’s lab. The lysosomes were captured on-grid using GFP nanobodies. These GFP nanobodies are attached to the maleimide groups covering the surface of the grid using a kck linker. All lysis steps were at 4°C and the lysate was kept on ice until plunge freezing.

Plunge freezing was done using a Leica GP2, a Whitman #1 blotting paper, and liquid ethane at -180°C as a cryogen for fast freezing. The functionalized grid was loaded into the chamber which was set at 4°C and 95% humidity. The lysate was pipetted up and down (20 strokes) in 10 ul volumes in 10 repeated rounds, resulting in an overall application of 100 ul of lysate to the grid surface. The grid surface was then washed with PBS two times by pipetting. Before adding the purified protein species most of the buffer volume left on the grid was pipetted away leaving the grid and the lysosomes attached to it just hydrated enough. This was done so that the dilution factor for the 6 purified protein species would be limited. Each of the purified proteins were added one at a time in 1 µl volumes as follows: 1) THG (from bovine thyroid, Sigma-Aldrich T9145) at a concentration of ∼17.8 mg/mL (A280=19.35), 2) Apoferritin (from equine spleen, Sigma-Aldrich 178440) at a concentration of ∼5 mg/mL (A280= 4.73), 3) Beta-galactosidase (from *E*. Coli, Sigma-Aldrich G5635) at a concentration of ∼6 mg/mL (A280=13.79), 4) Beta-amylase (from sweet potato, Sigma-Aldrich A8781) at a concentration of ∼5 mg/ml (A280=4.75), 5) Human Serum Albumin (Sigma-Aldrich A3782) at a concentration of ∼50 mg/mL (A280= 20.15), 6) Virus-like Particles (from collaborators at NYSBC) at a concentration of ∼7.5 mg/mL (A280=26.91). The target concentration for most of the species was aimed at 5 µM after the 6-fold dilution due to mixing. These are the per-species errors between the 5 µM goal and the final values as a result of challenges in volume handling at high concentrations: 1) THG: 15%, 2) Apoferritin: 33%, 3) Beta-galactosidase: 27%, 4) Beta-amylase: 16%. HSA concentration was kept high because it served as a background protein. VLP concentration could not be pushed further because of difficulties with volume handling at high concentrations. That said, all proteins are present in sufficient amounts in all the tomograms. After all the purified proteins were added to the grid, back-side blotting was done for 6 seconds and the grid was plunged into liquid ethane and then stored in liquid nitrogen.

### Data collection

All 1,089 tilt series were collected on one grid on a Krios G4 equipped with an X-FEG electron gun, a Falcon 4i direct electron detector, and the SelectrisX energy filter. The pixel size was set to 1.54 Å/pix, and the total dose to 62.93 e^-^/Å^2^ linearly spread over 31 tilt images spanning a range of -45° to +45° in 3° increments. The software used for data collection was TFS Tomo 5. Utilizing the beam-image shift data collection feature, 9 targets were imaged at each stage position. The movies were saved in the EER format.

For a rapid quality check of the sample, 2D data were also collected on the same grid for single particle analysis. The virus-like particles were refined to a resolution of 3.94Å. Briefly, 124 movie frames were collected and all the down-stream processing was done in CryoSparc^41^. To generate templates for VLPs, 12 micrographs were manually picked. Template matching resulted in 3,844 particles which were filtered down to 606 particles after 2D classification. Ab-initio model generation and the homogenous refinement resulted in a 3.93 Å map of an icosahedral VLP (Extended Data Fig. 8).

### Data processing

Motion correction, tilt-series alignment, and tomogram reconstruction were all performed using AreTomo3^26^. Specifically, raw frames were partitioned into non-overlapping groups of 2000 frames each, and batches of 10 frames within each group were integrated to generate rendered frames. Every two and four rendered frames were summed for measuring global and local motion, respectively. Motions measured on group sums were then interpolated to individual rendered frames for more accurate correction of rapid motion. For local motion estimates, these integrated frames were further subdivided into 5×5 patches. Both gain and motion corrected frames are summed to generate corrected tilt images. CTF parameters were estimated for each tilt image. These tilt images were then aligned into tilt-series, with global alignment followed by 4×4 patch-based local alignment. Tomograms were reconstructed using weighted-back projection, either with (for denoising) or without (for 3D template matching) applying a local CTF deconvolution and correction to the tilt-series. Even and odd pairs of CTF-deconvolved tomograms were produced to enable denoising. Paired tomograms were generated by splitting the motion-corrected frames into even and odd sets and then applying the alignment parameters determined for the full tilt-series to generate paired tomograms. Tomograms were denoised using denoisET, an in-house implementation of *Noise2Noise*^37^. This algorithm relies on a U-Net architecture composed of five downsampling and upsampling blocks each, with a kernel size of 3 pixels applied during each convolutional layer. This model was trained for 10 epochs on subvolumes extracted from 43 pairs of even and odd CTF-deconvolved tomograms before being applied to full tomograms to denoise the full dataset.

### Generating ground truth

Two approaches were taken to generate initial picks. For the 80S ribosome and VLP, PyTom’s^33,34^ 3D template matching algorithm was applied to the CTF-uncorrected and undenoised tomograms with a 10 Å pixel size. Templates were generated internally in PyTom from published maps (EMDB IDs: 3883 and 41917). For THG, apoferritin, and Beta-galactosidase, slab-picking was used to identify candidate particles. Specifically, denoised tomograms with a 5 Å pixel size were divided along the z-axis into non-overlapping 300 Å thick slabs, which were then projected along that axis to generate mock micrographs for further processing in CryoSPARC^41^. Blob picking was performed across a range of particle diameters, followed by iterative rounds of 2D classification to separate the different particle classes from each other and filter contamination based on visual inspection. Candidate particles were mapped back to their tomograms, and the depth of each particle was refined by locating the z-coordinate with the maximum integrated intensity in a subvolume centered on the particle.

Particle coordinates from PyTom and slab-picking were stored in copick, and candidate picks for 147 of the 492 tomograms were manually curated during a week-long pickathon marathon using the ChimeraX^35^ copick plug-in and copicklive to track curation statistics.

A more refined and expansive set of picks was generated using two deep learning approaches. One of these, DeepFindET (an adaptation of DeepFinder^12^), uses a Residual U-Net architecture with three 3D convolutional layers and a receptive field size of 680 Å to predict segmentation masks from annotated tomograms. The other, CellCanvas, involves a multi-step pipeline to segment tomograms. In the first step, a Swin UNETR model pre-trained on computed tomography data^42^ generates a voxel-wise embedding for each tomogram. In the second step, an interactively trained XGBoost classifier transforms clusters in embedding space into multi-class segmentation masks so that each voxel of the tomogram is assigned a probability for each class label. For initial training data, synthetic tomograms were generated in PolNet^43^ with a thickness of 180 nm and the five target particles, Beta-amylase, and membranes randomly distributed throughout the sample volume. An additive Gaussian noise model was applied, but neither the contrast transfer function or sample motion were modeled. The DeepFindET model was trained on 24 of these synthetic tomograms and then fine-tuned on five manually annotated tomograms from the phantom dataset and curated picks from the initial round of template matching and slab picking. For CellCanvas, the pre-trained Swin UNETR model was fine-tuned on four synthetic tomograms, while the classifier model was iteratively trained on manually annotated phantom tomograms. The segmentation masks predicted by DeepFindET were converted to individual picks using a size-based threshold, with the cut-off set to two thirds of each particle’s maximum dimension. In the case of CellCanvas, segmentation masks were grouped by the Watershed algorithm into similar regions, and the centroid of each region was labeled as the particle with the highest local probability density. For subsequent curation efforts, predictions for both models were pooled for THG and Beta-galactosidase, while the predictions for VLP, 80S ribosome, and apoferritin came exclusively from DeepFindET. We chose not to use the predictions from CellCanvas for these particles because the large number of candidate particles and particle miscentering proved computationally prohibitive to our downstream curation pipeline. Beta-amylase picks came from 2D classes of it that were generated during the classification of THG and Beta-galactosidase picks.

For the final rounds of curation, subvolumes centered on each particle were extracted from the denoised tomograms and projected along the z-axis to generate 2D per-particle projections. These “minislabs” were then tiled in 2D and loaded into CryoSPARC^41^ as mock micrographs, each containing 240 candidate particles. A single particle pick was positioned in the center of each tile. In the case of the VLPs, 80S ribosome, and apoferritin, we manually annotated 2, 14, and 19 micrographs per class respectively and used these positive labels to train a Topaz model^14^ for each particle. The trained Topaz models were then applied to the remaining micrographs. To increase the accuracy of our labels, we inspected minislabs of both the rejected particles to rescue any true positives and the accepted particles to eliminate any false positives during a final round of manual curation. In the case of the 80S ribosome, 2D classification followed by *ab initio* reconstruction with two classes was performed to assign a confidence score to each particle. Particles belonging to the manually-selected high-quality classes and retained in the correct reconstruction class were given a score of 1. Particles that segregated into the low-quality classes from 2D classification were assigned a score of 0, while the remaining particles that were selected during 2D classification but did not contribute to the correct reconstruction were assigned a score of 0.5. Only the score=1 ribosomes were retained for the challenge. In the case of THG and Beta-galactosidase, minislab micrographs were separately generated from the DeepFindET and CellCanvas picks, and 2D classification was performed on each set. After selecting high-quality classes based on visual inspection, the particles from each ML model were merged into a single set. Duplicates were removed based on a distance threshold of 185 Å and 125 Å for THG and Beta-galactosidase, respectively. The merged sets were manually curated, followed by *ab initio* reconstruction with two classes.

Particles that contributed to the correct reconstruction class were retained. Finally, we generated minislabs from any positions that belonged to more than one particle class and manually assigned one label based on visual inspection to remove these inter-class duplicates. The final particle lists from these curation efforts were considered the ground truth annotations.

## Software Availability

Software packages used in this study for reconstruction of tomograms, annotation generation and annotation curation are released under open source licenses and available in public repositories. Refer to **Supplementary Table 1** for a summary of packages.

## Contributions

**M.P.**, **K.H.,** and **B.C.** developed the idea for a machine learning challenge and managed the project. **M.P.**, **H.S.**, **D.S.**, and **F.W.** optimized grid functionalization and freezing of organelles and carried out the sample preparation. **M.P.** and **E.M.** carried out the data collection. **M.P.** curated the tomograms based on sample quality and tilt series alignment quality. **S.Z.**, **A.P.**, **Y.Y.**, **J.S.**, and **M.P.** developed the data processing pipeline. **S.Z.** and **A.P.** developed AreTomo3 CTF deconvolution and DenoisET. **S.Z.**, **A.P.**, and **Y.Y.** developed the slab method. **U.H.E.** developed copick and ChimeraX-copick plugin. **K.H.** and **Z.Z.** developed CellCanvas and copicklive. **J.S.** developed DeepFindET. **D.K.** and **J.S.** developed the 3D refinement pipelines. **K.H.**, **Z.Z.**, and **J.S.** developed the challenge metrics and the dataset splitting. **M.P.** and **Y.Y.** manually picked initial training sets. **M.P.**, **A.P.**, **Y.Y.**, and **J.S.** prepared the final curations for all the picks for all the species. **S.K.**, **J.S.**, **K.H.**, and **Z.Z.** prepared the example notebooks. **U.H.E.** and **A.C.** developed the cryoET data portal. **M.P.**, **A.P.**, **U.H.E.**, **K.H.**, **Z.Z.**, and **B.C.** wrote the manuscript. All authors reviewed the manuscript. **M.H., D.A., C.P.** and **B.C**. provide overall leadership for projects at the Chan Zuckerberg Imaging Institute.

## Acknowledgements

- These authors contributed equally to this work and were listed alphabetically: Anchi Cheng, Utz Heinrich Ermel, Saugat Kandel, Dari Kimanius, Elizabeth Montabana, Daniel Serwas, Hannah Siems, Feng Wang, Zhuowen Zhao, Shawn Zheng.
- Emma Lundberg (Stanford), Ellen Zhong (Princeton), Thorsten Wagner (MPI of Molecular Physiology), Tristan Bepler (NYSBC), Robert Kiewisz (NYSBC), Alister Burt (Genentech), and Lorenzo Gaifas (Grenoble) who provided valuable insights for the design of the challenge.
- Our collaborators at CZ BioHub SF, Manuel Leonetti, Shivanshi Vaid, Madhuri Vangipuram, and Rodrigo Baltazar generated the HEK293T LAMP1-GFP cell lines and shared their cell lysis protocol.
- David Agard’s lab at UCSF, Feng Wang and Simon Sander, provided their grid functionalization protocol and grids and helped us optimize the protocol further for organelles.
- The GFP-nanobody construct was modified and provided by Peng Jin (UCSF, Jan Lab).
- Our collaborators at NYSBC, Mykhailo Kopylov and Charlie Dubbledam, provided VLPs.
- Chan Zuckerberg Initiative SciTech members who participated in the pickathon: Ashley Anderson, Ben Nelson, Jun Ni, Ellaine Chou, Jessica Gadling, Kandarp Khandwala, Chili Chiu, Ann Jones, Timmy Huang, Janeece Pourroy, Dannielle McCarthy, Andy Sweet, Eric Wang, Kirsty Ewing, Mikala Caton, Manasa Venkatakrishnan, Kira Evans.
- CZ Imaging Institute members who participated in the pickathon: Yongbaek Cho, Nina Borja, Norbert Hill, Carmela Villegas, Shu-Hsien Sheu, Gorica Margulis, Noeli Pazsoldan.
- Chan Zuckerberg Initiative SciTech team who contributed to the development of the cryoET data portal: Jun Xi Ni, Jessica Gadling, Manasa Venkatakrishnan, Kira Evans, Jeremy Asuncion, Andrew Sweet, Janeece Pourroy, Zun Shi Wang, Kandarp Khandwala, Benjamin Nelson, Dannielle McCarthy, Eric M Wang, Richa Agarwal, Trent Smith, Bryan Chu, Dana Sadgat, Erin Hoops, Justine Larsen.
- Kristen Maitland and Stephani Otte for their support in the planning and execution of the competition.
- Samantha Yammine who provided valuable feedback on the manuscript text.
- Some schematic elements in Figure 1 (panels a,b), Figure 2 (panels a,b,c) and Extended Data Figure 1 were made using BioRender: Created in BioRender. Paraan, R., Serwas, D. (2024) BioRender.com/r59u258
- Grant Reference: CZ Imaging Institute is made possible with support from Chan Zuckerberg Initiative (CZII-2023–327779).

