## Supplemental figures and tables for "Annotating CryoET Volumes: A Machine Learning Challenge"

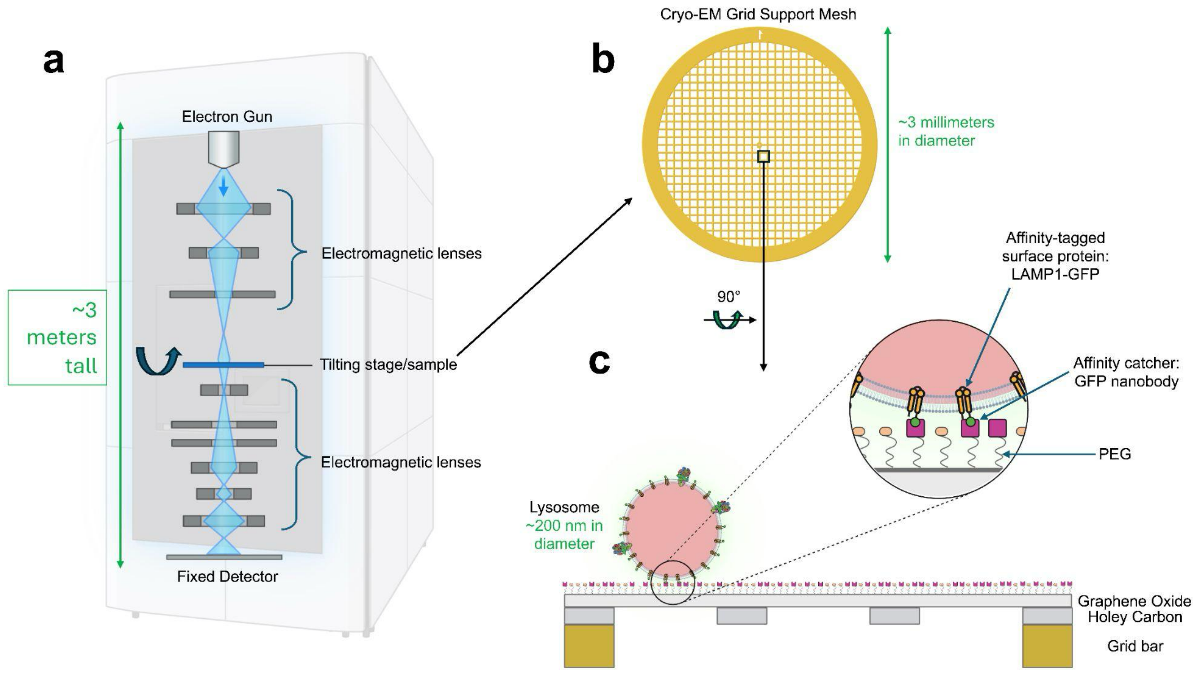


**Extended Data Figure 1 | Overview of the microscope and sample technologies used in this study. a**. Simple schematics of the microscope column showing the electron gun at the top, followed by condenser electromagnetic lenses used for focusing a parallel coherent beam on the sample, followed by the compustage that holds and rotates the sample for tomography data collection, followed by lenses that converge and magnify the beam on the detector. There are important pieces, such as the energy filter, that are not shown for simplicity. **b**. The platform or the so-called electron microscopy grid that holds the sample is made of a mesh structure, in this case made of gold, that supports a thin carbon film with repetitive holes (not shown). **c**. 90° cross-section view of one of the squares from the mesh of a functionalized grid. The different layers and structures are not to scale for illustration purposes. The process of functionalization involves adding the graphene oxide layer, the PEG layer, and the GFP nanobodies. These GFP nanobodies will bind LAMP1-GFP on the surface of lysosomes present in cell lysates.

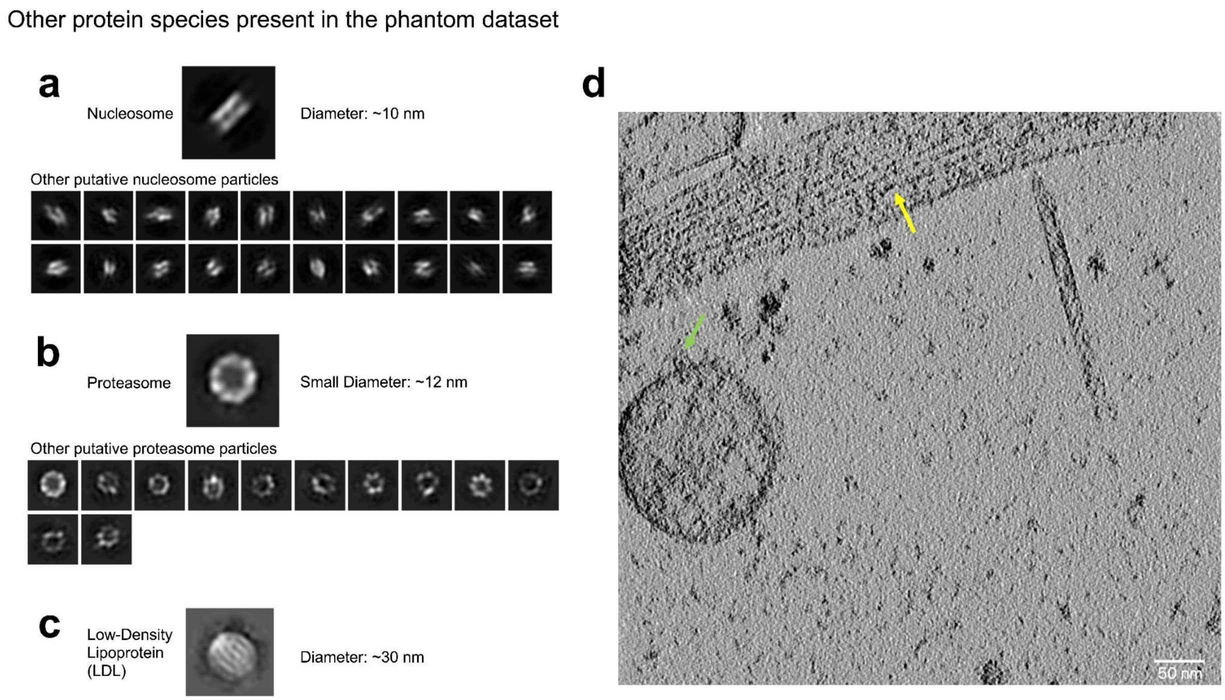


**Extended Data Figure 2 | Other species of proteins that co-purified from the lysate and are NOT part of the challenge**. **a**. 2D classes of nucleosomes. They are prevalent in the nucleus and co-purify after cell lysis. **b**. 2D classes of proteasomes. They are cytosolic and less prevalent than nucleosomes in the dataset. Their small diameter as seen in the 2D class is ~12 nm and their large diameter (not shown) is ~16 nm. **c**. A 2D class of LDL complexes. They are less prevalent than nucleosomes and proteasomes. They attach to membranes. **d**. An example tomogram slice showing a bundle of actin filaments (yellow arrow) and transmembrane protein complexes (lysosomal proteins, green arrow). These are scattered randomly across the dataset and are not highly abundant.

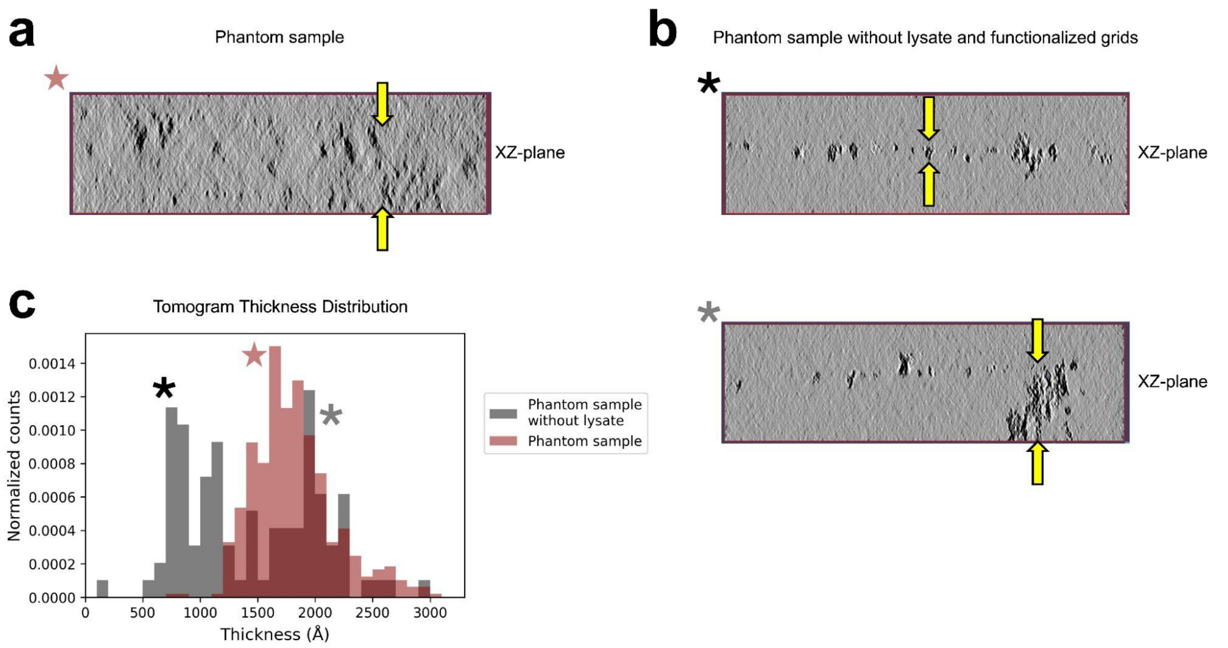


**Extended Data Figure 3 | Graphene-oxide functionalized grids and cell lysates create more uniform and thicker samples**. **a**. XZ view of the phantom dataset. Yellow arrows point to the boundaries of the sample and indicate a thicker volume compared to b. This thicker volume captures the typical noise behavior in a cryoET dataset. **b**. XZ views of another sample prepared in-house which is a mix of 5 proteins (VLP, apoferritin, BSA, THG, Beta-amylase). Without using graphene-oxide functionalized grids and lysates, the sample is mostly one layer of proteins (top). This introduces issues like aggregation (bottom) and SNR similar to single particle analysis as opposed to cryoET. **c.** The thickness of the tomograms from the two datasets were estimated using Aretomo3[26]. The phantom dataset (a) shows a uniform distribution. The phantom dataset without cell lysates and functionalized grids (b) has a bimodal distribution: one peak (around 600 Å) for tomograms that are one particle-layer thick (b, top), and another peak (around 2000 Å) for tomograms with large aggregations (b, bottom).

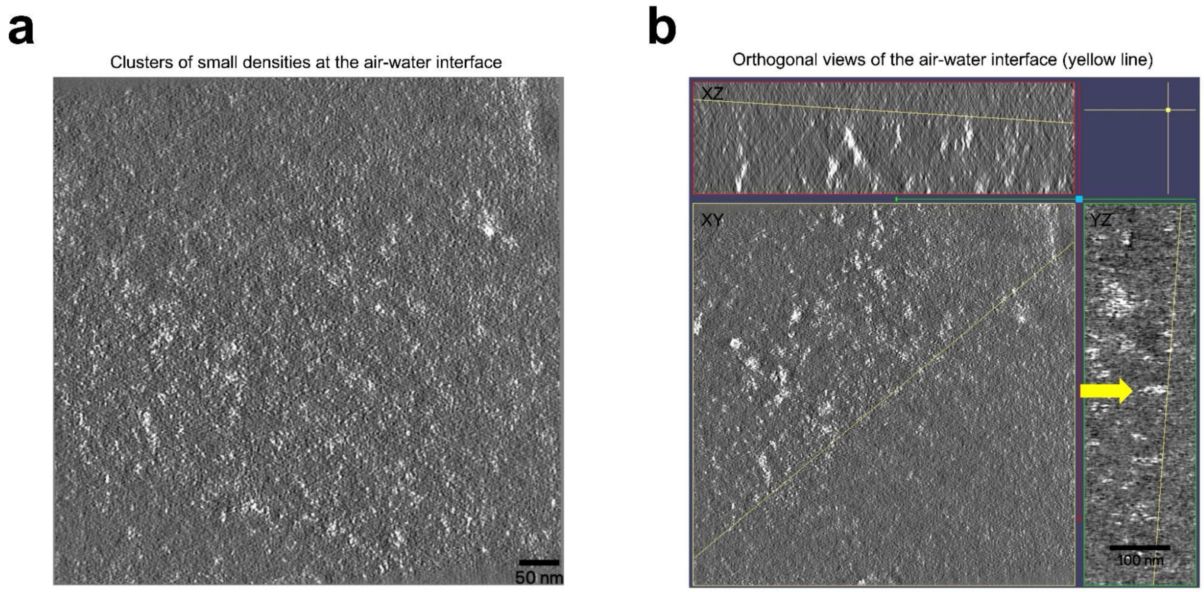


**Extended Data Figure 4 | The air-water interface in the phantom dataset. a**. A slice through the air-water interface showing a large population of small densities as white speckles. Scale bar is 50 nm. **b**. The same tomogram from a, seen in orthogonal views. The yellow line indicates the common line between the tilted plane that generated panel a and each orthogonal view. The yellow arrow shows a larger density at the air-water interface. The contrast was inverted to white particles and dark background for better visualization of the small densities. Scale bar is 100 nm.

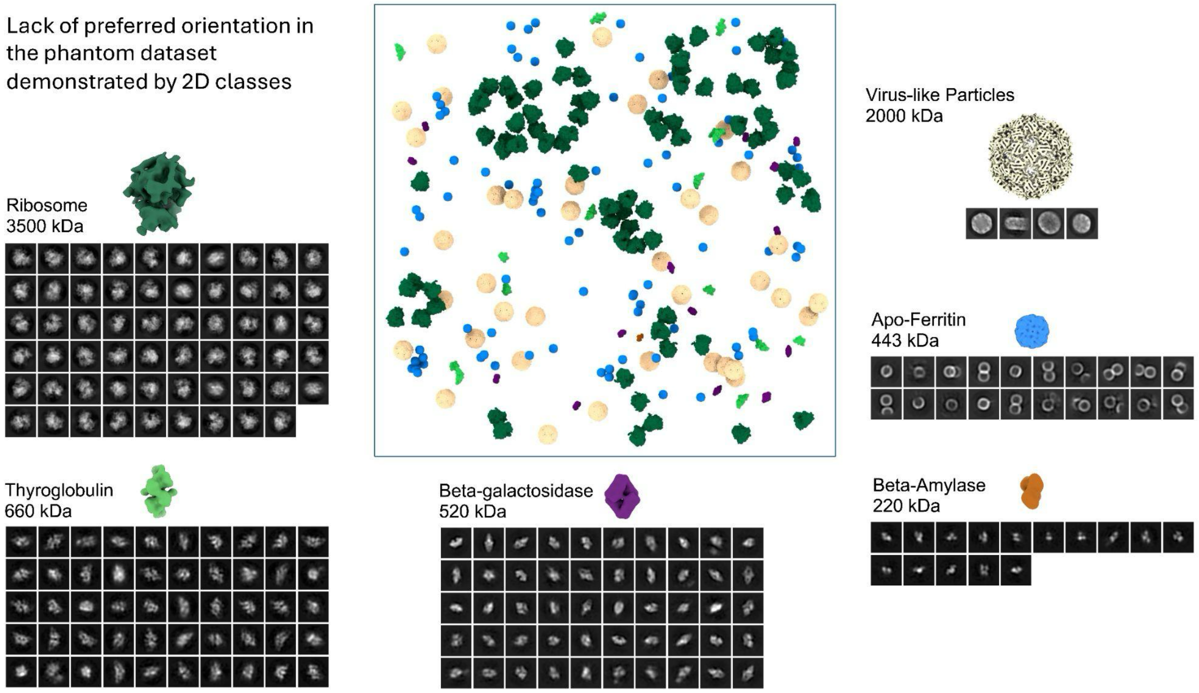


**Extended Data Figure 5 | Lack of preferred orientation in the phantom dataset.** 2D classes were generated using the slab method on denoised tomograms with a pixel size of 5 Å/pix. For ribosome, thyroglobulin, Beta-galactosidase, VLP, and Beta-amylase species, the 3D maps were generated from the phantom data using the slab method (higher resolution maps are currently being processed and will be released soon). For apoferritin, the 2d classes are generated from the data, but the 3D map is a representative published map. This is because the apoferritin in this sample is not optimized for high-resolution structure determination, as is typically required for apoferritin. VLP and apoferritin are highly symmetric, therefore, the 2D classes are representative of conformational heterogeneity rather than the orientation distribution.

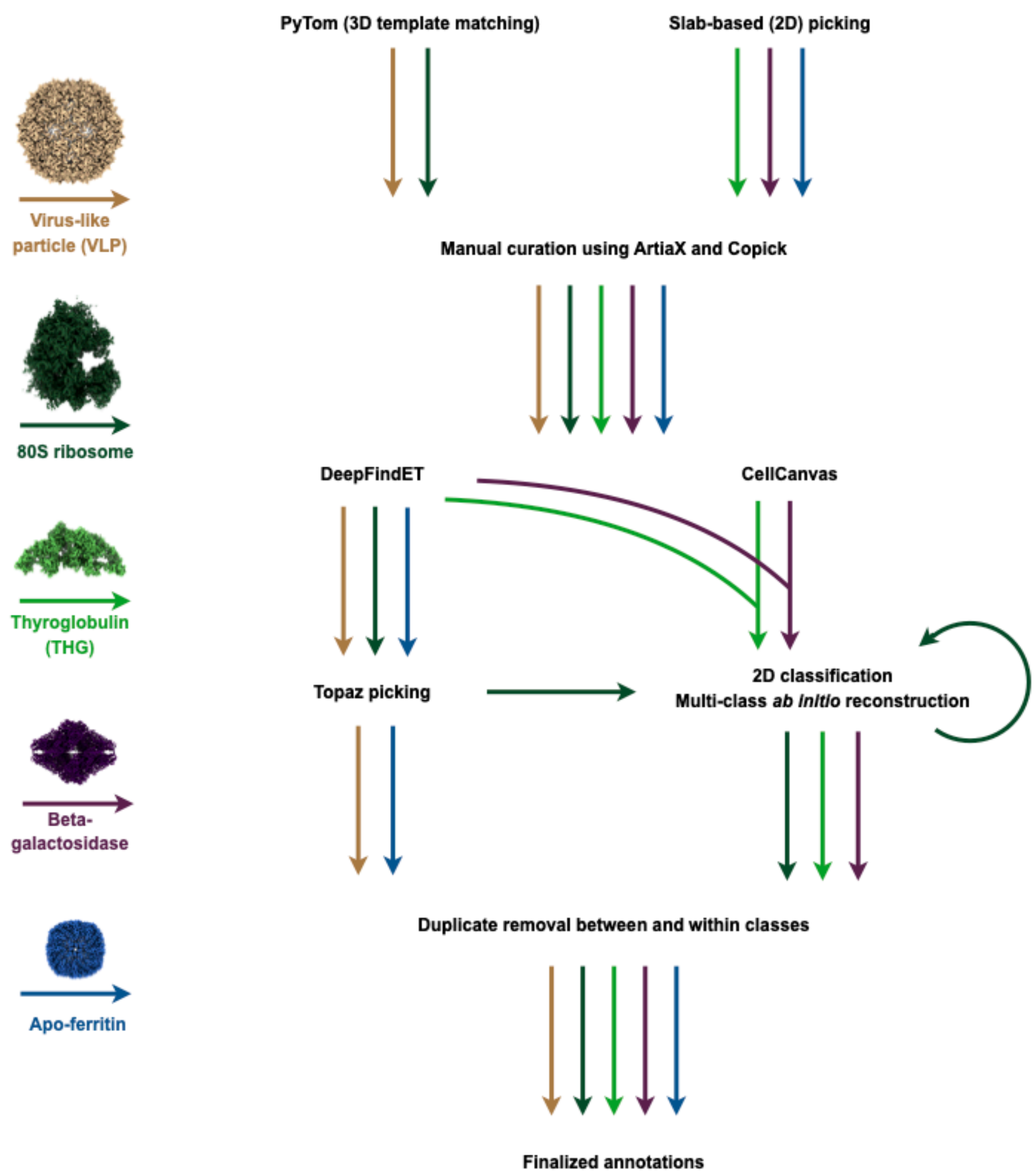


**Extended Data Figure 6 | Workflow for generating ground truth labels.** Despite applying each of the listed picking and refinement methods to every particle, we found that no single workflow worked for all particles. This schematic traces which tools were applied to generate the final annotations that we refer to as ground truth labels for each particle class, with each particle indicated by the color shown in the legend at the left.

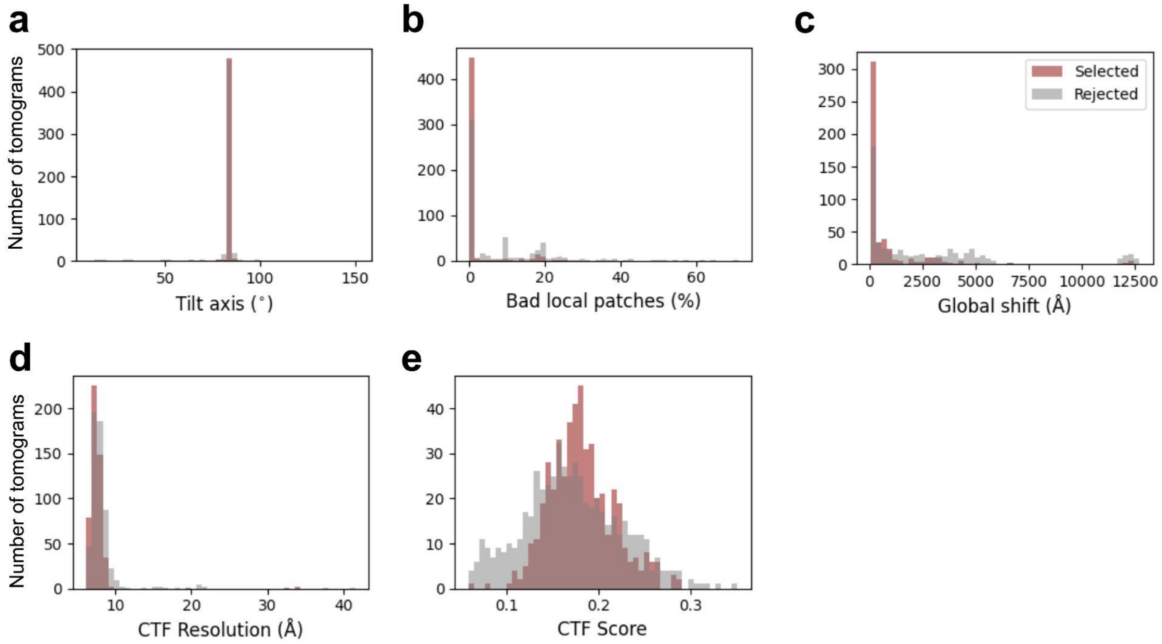


**Extended Data Figure 7 | AreTomo3 tilt series quality metrics for the phantom dataset (1,073 tomograms total) plotted against the tomograms that were manually curated for quality (492 tomograms selected). a**. Selected tomograms fall within the correct tilt axis angle. **b**. Selected tomograms have the lowest percentage of bad patches in patch-based tilt series alignment. **c**. Selected tomograms have the lowest global shifts during data collection. **d**. Selected tomograms have average CTF resolutions for tilt series that are below 10 Å. **e**. Selected tomograms have the highest CTF confidence scores.

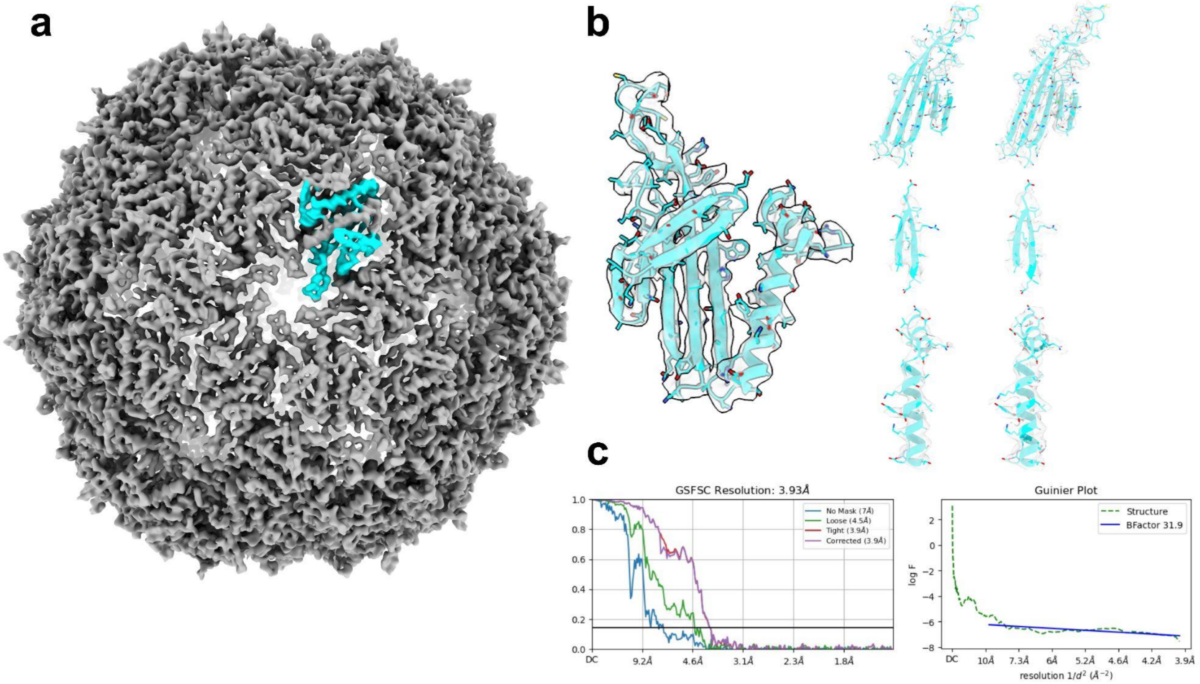


**Extended Data Figure 8 | High-resolution reconstruction of icosahedral VLP from 2D micrographs (single particle analysis) from the same phantom sample**. **a**. Icosahedral map of VLP refined in CryoSparc[41] shows clear side chain densities. The asymmetric subunit is highlighted in cyan and expanded in panel b. **b**. The atomic model of the subunit (PDB: 1DWN) is rigidly fitted into the subunit density. A full subunit is shown on the left side, and different secondary structures of the same subunit are shown on the right. For each part of the subunit on the right, the map is drawn at two different isosurface thresholds. The lower threshold on the right shows side chain densities for those that are more sensitive to radiation damage. **c**. The FSC plot on the left indicates a resolution of 3.93 Å. The B-factor plot on the right indicates a B-factor of 31.9 Å$^2$, which indicates a high map quality[54].

### Supplemental Tables

| Software Package | Repository Link | License |
| --- | --- | --- |
| Copick | https://github.com/copick/copick | MIT |
| ChimeraX-Copick | https://github.com/copick/chimerax-copick | MIT |
| Copicklive | https://github.com/copick/copick_live | MIT |
| CellCanvas | https://github.com/cellcanvas/cellcanvas | MIT |
| AreTomo3 | https://github.com/czimaginginstitute/AreTomo3 | BSD 3-Clause |
| DeepFindET | https://github.com/copick/DeepFindET | GPLv3 |
| Slabpick[a] | https://github.com/apeck12/slabpick | MIT |

[a] *slabpick* was used for both the slab-picking and minislab-curation workflows.

**Supplementary Table 1: Software Availability**. Source code, open-source licenses and references (if applicable) for tools used for tomogram reconstruction, annotation generation and annotation curation in this study.

| Notebook | Description | Link |
|---|---|---|
| 3D U-Net | A 3D U-Net built on MONAI that uses copick projects for training and prediction | https://github.com/czimaginginstitute/2024_czii_mlchallenge_notebooks/tree/main/3d_unet_monai |
| TomoTwin | An application of the generalist particle picking tool, TomoTwin, for inference. | https://github.com/czimaginginstitute/2024_czii_mlchallenge_notebooks/tree/main/tomotwin_picking_notebook |
| DeepFindET | A ResUNet-based model derived from DeepFinder that uses copick projects for trianing and prediction. | https://github.com/czimaginginstitute/2024_czii_mlchallenge_notebooks/tree/main/DeepFindET |

**Supplementary Table 2: Example notebooks**. These example notebooks are based on three published models and they are designed such that cryoET domain non-experts can get a headstart. These notebooks provide a full workflow for training, predicting, and submitting to Kaggle and they are meant to familiarize the participants with the copick libraries and metadata/data handling.
